# Targeting c-Jun is a potential therapy of luminal breast cancer bone metastasis

**DOI:** 10.1101/2022.07.17.500330

**Authors:** Yuxuan Han, Mitsuru Futakuchi, Kazuya Nakamichi, Yutaro Wakabayashi, Mai Sakamoto, Jun Nakayama, Kentaro Semba

## Abstract

Luminal breast cancer has the highest bone metastasis frequency among all breast cancer subtypes, but its metastatic mechanism has not been elucidated because of the lack of appropriate metastatic cell lines. The study aim was to characterize high-osteolytic bone metastatic MCF7-BM cell lines and extract c-Jun, a novel bone metastasis marker. We found that c-Jun was upregulated in MCF7-BM cells, and its deficiency was associated with suppression of the cell migration, transformation, and stemness of BM cells. *In vivo*, c-Jun-deficient MCF7-TAM67 cells exhibited weaker bone metastatic ability. Additionally, c-Jun overexpression in MCF7-BM cells led to a tumor-migration promotion cycle in the bone microenvironment possibly by enhancing calcium-induced migration and releasing the osteoclast activator BMP5. Inhibition of c-Jun by JNK-IN-8, a JNK inhibitor, effectively reduced tumorigenesis activities and bone metastatic tumors. Our results indicate the potential benefits of a therapy that targets c-Jun to prevent or minimize luminal breast cancer bone metastasis.

## Introduction

Bone is the major metastatic site of breast cancer^1^. Although bone metastasis is not fatal, it is generally incurable by available clinical treatments and causes a series of complications called skeletal-related events, which have a poor prognosis^2^ and directly affect the quality of life^3,4^. Bone metastasis frequency varies among breast cancer subtypes. The bone metastasis frequency of the luminal subtype is up to twice as high as that of other subtypes^5,6^. In contrast, triple-negative breast cancer (TNBC) tumors have a higher rate of brain, liver, and lung metastases but a lower rate of bone metastasis than luminal tumors^7^. Nevertheless, most breast cancer metastasis studies have been performed using MDA-MB-231, a TNBC breast cancer cell line^8,9^. However, no such suitable cell model has been established for the luminal subtype so far^10^. We previously established highly bone metastatic MCF7 (MCF7-BM) cell lines and analyzed their application in a study of luminal breast cancer bone metastasis^10^. MCF7-BM cells use a distinct mechanism from that of MDA-MB-231 cells to activate osteoclasts and acquire osteolytic bone metastasis. Our previous study highlighted the need for a subtype-specific study on bone metastasis of breast cancer^11^.

The AP-1 transcription factor, c-Jun, was originally discovered as an oncogene of avian sarcoma virus^12^ and is known to have a critical role in the invasion and metastasis of human breast cancer cell lines. Overexpression of c-Jun in MCF7 cells promotes tumorigenic and hormone-resistant ability^13^. By regulating stem cell factor and C–C chemokine ligand 5, c-Jun can induce mammary epithelial cellular invasion and breast cancer stem cell expansion^14^. A xenograft model showed that c-Jun overexpression was also relevant to the increased liver metastasis of SKBR3 cells, a HER2-positive breast cancer cell line^15^. Silencing of c-Jun can decrease cellular migration and invasion in CNE-2R, a radio-resistant human nasopharyngeal carcinoma cell line^16^. Additionally, c-Jun is expressed in primary and metastatic lung tumors in 31% of cases^17^. Although many studies have focused on the function of c-Jun during tumor progression, its role in bone metastasis and the underlying mechanism have not been investigated in detail.

In this study, we analyzed the function of c-Jun overexpression in MCF7-BM cells and evaluated the potential of targeting c-Jun as a therapy against bone metastasis of luminal-type breast cancer.

## Results

### Involvement of c-Jun in malignant transformation and osteolytic bone metastasis of MCF7 cells

We previously established high-osteolytic luminal bone metastatic cell lines, BM01 and BM02, from MCF7 cells^10^. In the present study, we investigated the difference in signal transducers between these metastatic cell lines and the parental cell line by western blotting. We found that the protein levels of c-Jun and phosphorylated c-Jun at Ser73 were increased, whereas the phosphorylation level of c-Fos, a binding partner of c-Jun to constitute AP-1 factor, did not show any difference. Additionally, the phosphorylation level of Jun amino terminal kinase (JNK) showed no change in BM cells (Fig. 1a). This result suggested that the increased p-c-Jun expression was due to an increased protein level of c-Jun. Both the c-Jun and p-c-Jun proteins are almost exclusively localized in the nucleus (Fig. 1b), suggesting aberrant gene expression by increased c-Jun. The mRNA level of c-Jun exhibited no significant difference between parental cells and BM02 cells, indicating that the increase in c-Jun was caused by a post-transcriptional mechanism (Fig. 1c).

**Fig. 1.**
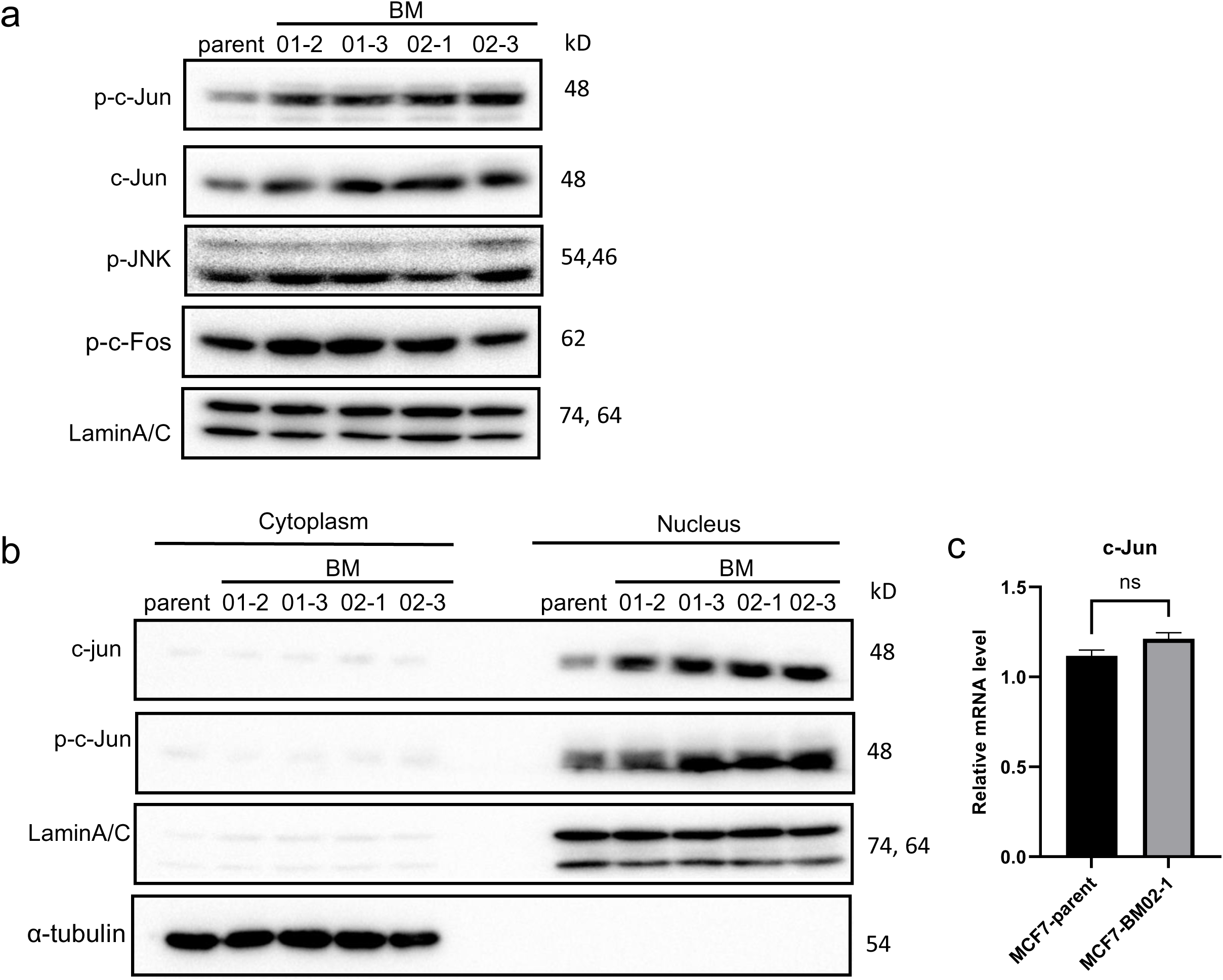
c-Jun protein was overexpressed in MCF7-BM cells. **a)** The cell lysates of the MCF7-parent, MCF7-BM01-2, MCF7-BM01-3, MCF7-BM02-1, and MCF7-BM02-3 were collected and subjected to western blotting. LaminA/C was the loading control. **b**) The nucleus and cytoplasm lysates of the MCF7 cell lines were analyzed separately to localize the c-Jun and p-c-Jun proteins. LaminA/C and α-tubulin were used as the loading control. **c)** The mRNA levels of c-Jun in the MCF7-parent and MCF7-BM02-1. The data are presented as the mean ± SEM. The statistical analysis was performed by applying the *t*-test with Welch’s analysis.

To investigate the function of c-Jun overexpression in MCF7 cells, we established a c-Jun overexpressed MCF7-parent cell line, MCF7-JUN (Fig. 2a). Overexpression of c-Jun did not affect the cell proliferation ability of MCF7 *in vitro* (Fig. 2b). MCF7-JUN showed enhanced wound-healing (Fig. 2c, d), colony formation (Fig. 2e, f), and tumorsphere ability (Fig. 2g, h), which suggested that c-Jun overexpression enhanced cell migration, transforming ability, and tumor stemness. We next performed a Tartrate-resistant acid phosphatase (TRAcP) co-culture assay between osteoclasts and MCF7 cells to evaluate the function of overexpressed c-Jun toward osteoclast activation. TRAcP-positive osteoclasts were significantly increased under co-culture with MCF7-JUN, indicating that c-Jun overexpression in MCF7 cells could activate osteoclasts and promote their maturation (Fig. 2i, j).

**Fig. 2.**
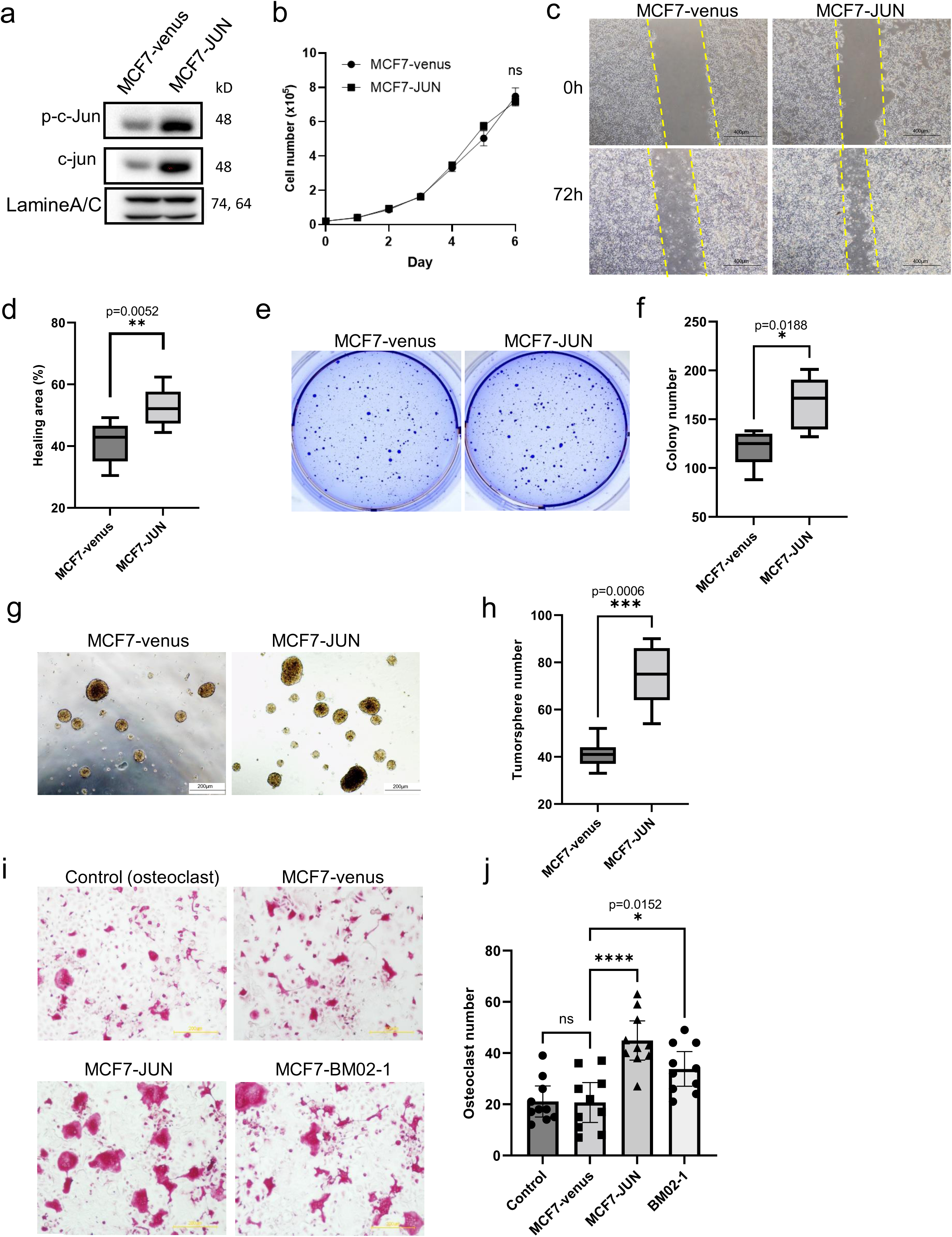
Characterization of the c-Jun overexpressed MCF7-JUN cell line. **a)** A c-Jun-overexpressed cell line MCF7-JUN was established from the MCF7-parent cell line. The c-Jun protein expression was confirmed by western blotting. MCF7-venus was established as a transfection control. **b)** The growth curve of MCF7-venus and MCF7-JUN cells. The data are presented as the mean ± SEM. The statistical analysis was performed by applying the *t*-test with the Mann–Whitney test. **c, d)** The scratch assay of the MCF7-JUN cell line. **c** Photos of the wound-healing process. **d** Statistical analysis of the healing area. The results were summarized from two independent experiments and analyzed by the *t*-test with Mann–Whitney analysis. **e, f)** The soft agar-formation assay of the MCF7-JUN cell line. **e** Photos of crystal violet staining of colonies. **f** Statistical analysis of the colony number. Summary of the results from two independent experiments analyzed by the *t*-test with Mann–Whitney analysis. **g, h)** The tumorsphere assay of MCF7-JUN. **g** Photos of tumorspheres at the 7^th^ day of culture. **h** Statistical analysis of the number of tumorspheres. The results were summarized from two independent experiments and analyzed by the *t*-test with Mann–Whitney analysis. **i, j)** The tartrate-resistant acid phosphatase (TRAcP) MCF7-JUN cells and osteoclasts co-culture assay. **i** Photos of mature osteoclasts stained by TRAcP; the controls were the osteoclasts without tumor-cell interaction. **j** Statistical analysis of the TRAcP assay results. The results were summarized from two independent experiments and analyzed by one-way ANOVA with Dunnet’s test.

Next, we examined the suppression of c-Jun function in BM02 cells by introducing the dominant negative c-Jun, TAM67^18^. Since c-Jun transcription is positively autoregulated by its own product, Jun^19^, BM02 expressing the TAM67 (BM02-TAM67) cell line showed decreased c-Jun expression (Fig. 3a, b). Introduction of TAM67 did not affect the cell proliferation ability of BM02 (Fig. 3c). The TAM67 expression significantly reduced cell migration (Fig. 3d, e), transforming ability (Fig. 3f, g, h), and tumor stemness (Fig. 3i, j) of BM02. Dominant negative c-Jun also suppressed the osteolytic ability of BM02 cells (Fig. 3k, l).

**Fig. 3.**
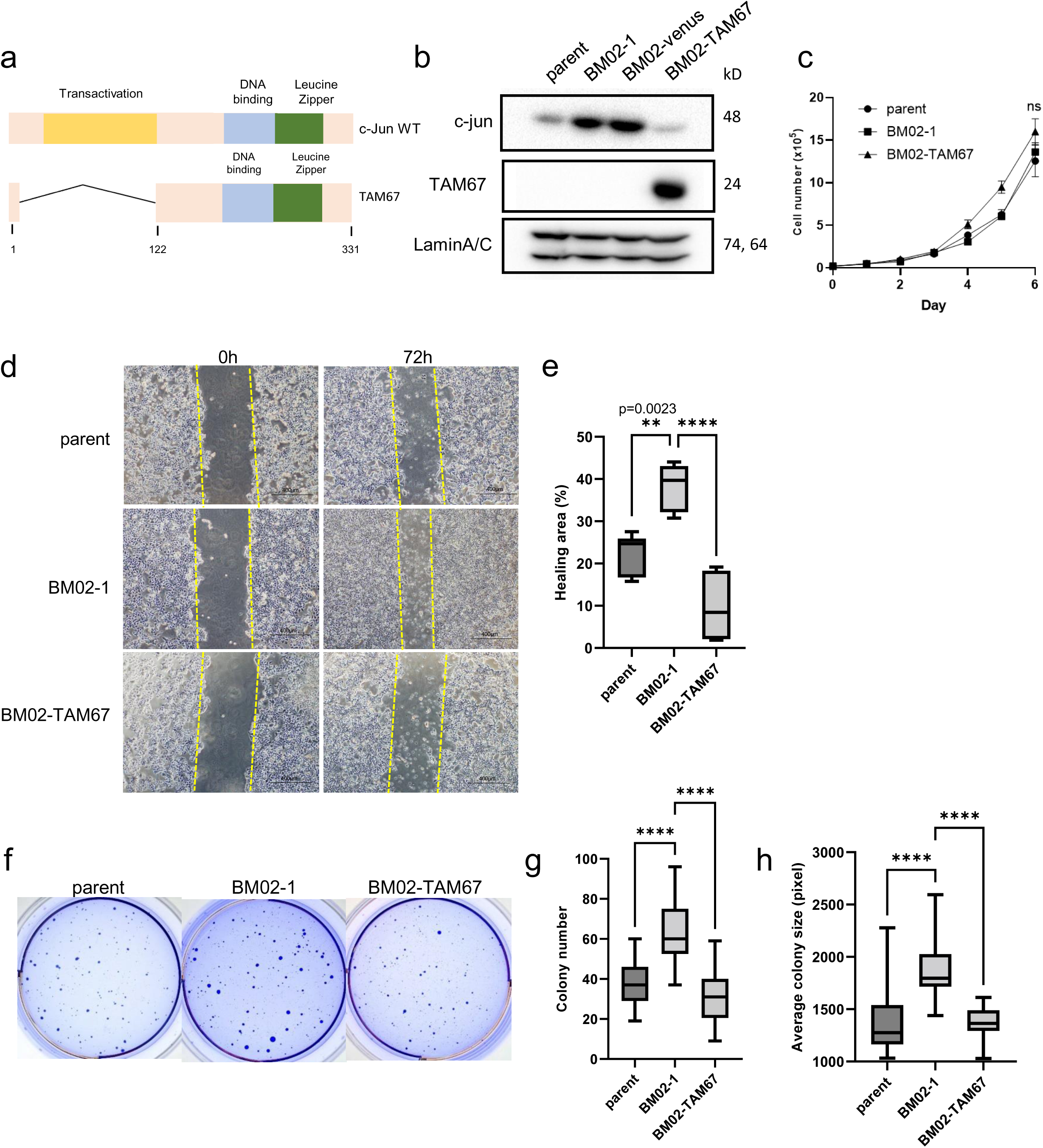
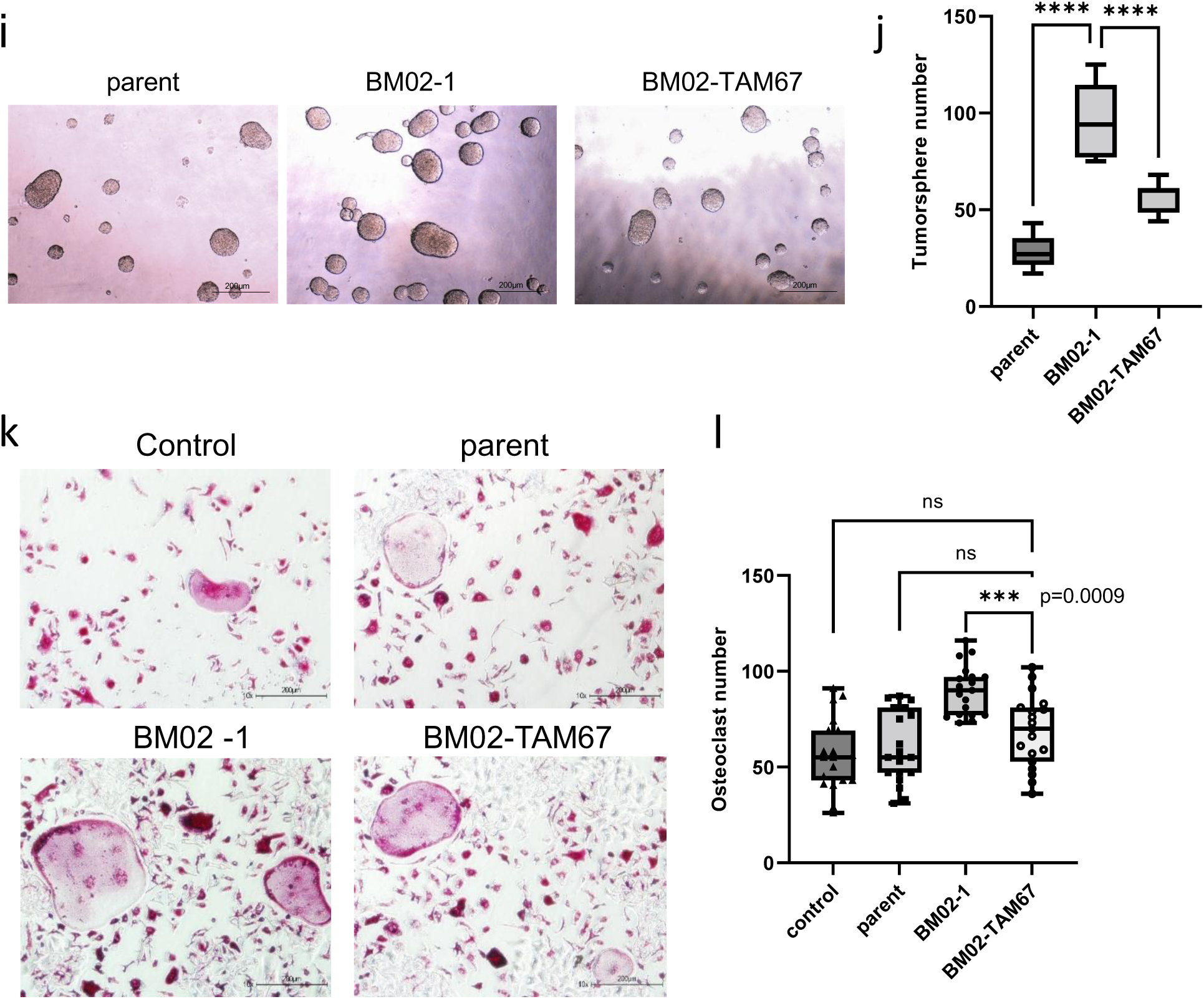
Establishment and characterization of the c-Jun-deficient BM02-TAM67 cell line. **a)** A scheme showing TAM67 and c-Jun structure. **b)** A c-Jun-deficient cell line, BM02-TAM67, was established from the MCF7-BM02-1 cell line. The TAM67 protein expression was confirmed by western blotting. MCF7-BM02-venus was established from MCF7-BM02-1 as a transfection control. **c)** Growth curve of the BM02-TAM67 cell line. The data are presented as the mean ± SEM and analyzed by *t*-test with the Mann–Whitney analysis. **d, e)** Scratch assay of the BM02-TAM67 cell line. **d** Photos of the wound-healing process. **e** Statistical analysis of the healing area. The results were summarized from two independent experiments and analyzed by one-way ANOVA with Dunnet’s test. **f, g, h)** The soft agar-formation assay of the BM02-TAM67 cell line. **f** Photos of crystal violet staining of colonies. **g** Statistical analysis of colony number. **h** Statistical analysis of the average colony size by pixel. The results were summarized from three independent experiments and analyzed by one-way ANOVA with Dunnet’s test. **i, j)** The tumorsphere assay of BM02-TAM67. **i** Photos of the tumorspheres at the 7^th^ day of culture. **j** Statistical analysis of the number of tumorspheres. The results were summarized from three independent experiments and analyzed by one-way ANOVA with Dunnet’s test. **k, l)** The TRAcP co-culture assay between BM02-TAM67 cells and osteoclasts. **k** Photos of mature osteoclasts stained by TRAcP; the controls were the osteoclasts without tumor-cell interaction. **l** Statistical analysis of TRAcP assay. The results were summarized from three independent experiments and analyzed by one-way ANOVA with Dunnet’s test.

On the basis of the above results, we transplanted BM02-TAM67 by intra-caudal arterial injection (CAI) into NOD/scid mice (Fig. 4a, b, Supplementary Fig. S1a). Compared with BM02, BM02-TAM67 had smaller bone metastatic lesions (Fig. 4c, d) and a lower bone metastasis frequency (Fig. 4e). Histological analysis showed that the BM02-TAM67 bone lesions exhibited non-osteolytic metastasis (Fig. 4f), and TAM67 cells failed to activate osteoclasts within the bone microenvironment (Fig. 4g). The expression of ki67 was overall decreased in BM02-TAM67 relative to that in BM02 (Fig. 4h, i), suggesting that BM02-TAM67 was a less progressive cell line.

**Fig. 4.**
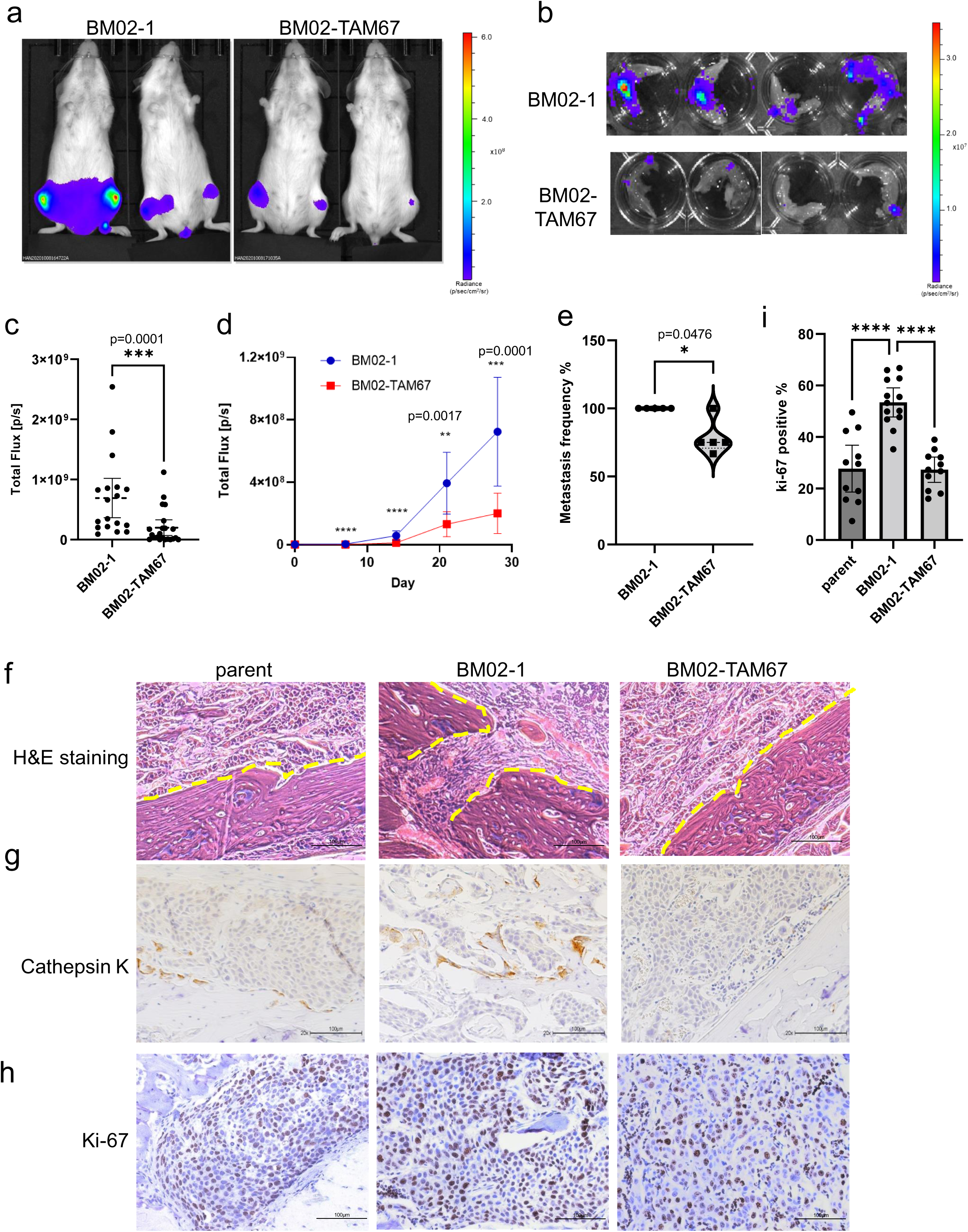
c-Jun deficiency reduced tumorigenesis activity and bone metastatic tumor. **a, b)** Transplantation of BM02-1 and BM02-TAM67 to NOD/scid mouse through intra-caudal arterial injection (CAI). The transplantation was performed independently five times (BM02-1, n=9; BM02-TAM67, n=11). **a** *In vivo* imaging photos of a mouse with bone metastatic lesions. **b** *Ex vivo* imaging photos of legs with bone metastatic tumors. **c, d, e)** Statistical analysis of CAI. The data are presented as the mean ± 95%CI and analyzed by one-way ANOVA with Dunnet’s test. **c** Summary of tumor size at day 28. **d** Tumor growth through *in vivo* observation. **E** Summary of the metastatic frequency from five CAI transplantations. Paraffin sections of metastatic lesions stained with **f)** hematoxylin and eosin (H&E) staining; the yellow line indicates the bone surface; **g)** Anti-cathepsin K (osteoclast marker) and **h)** anti-ki-67 (tumor proliferation marker), and observed under an optical microscope at x20 magnification. **i)** Statistical summary of ki-67 histological expression. The data are presented as the mean ± 95%CI. The significance was calculated by one-way ANOVA with Dunnet’s test.

### Possible vicious cycle induced by c-Jun in the bone microenvironment

Histological analysis was also used to evaluate c-Jun expression (Fig. 5a, b) and showed that c-Jun expression in BM02 was highly heterogenous. Tumor cells with high levels of c-Jun tended to gather around the destroyed bone surface, whereas the tumor cells near the non-destroyed bone surface exhibited relatively low c-Jun expression (Fig. 5c, Supplementary Fig. S1b). This finding suggested that tumor cells with high expression of c-Jun were able to interact with factors released from the destroyed bone surface and further promoted tumor-migration. A previous study found that various factors that contribute to the invasion and proliferation of tumor cells were released from destroyed bone surfaces ^20^. Among these factors, we noticed that in contrast to the trans-well results of MCF7 cell lines in FBS (Fig. 5d, e), more BM02 cells and MCF7-JUN cells migrated to a high Ca^2+^ environment than parental cells, whereas BM02-TAM67 cells lost that ability (Fig. 5d, f). These results suggested that high expression of c-Jun in MCF7 cells induced cell migration towards high Ca^2+^ environment. This ability of BM02 cells may further contribute to the interaction between osteoclasts and tumor cells. Consistently, Ca^2+^ concentration is known to be higher around the bone resorption area^21,22^.

**Fig. 5.**
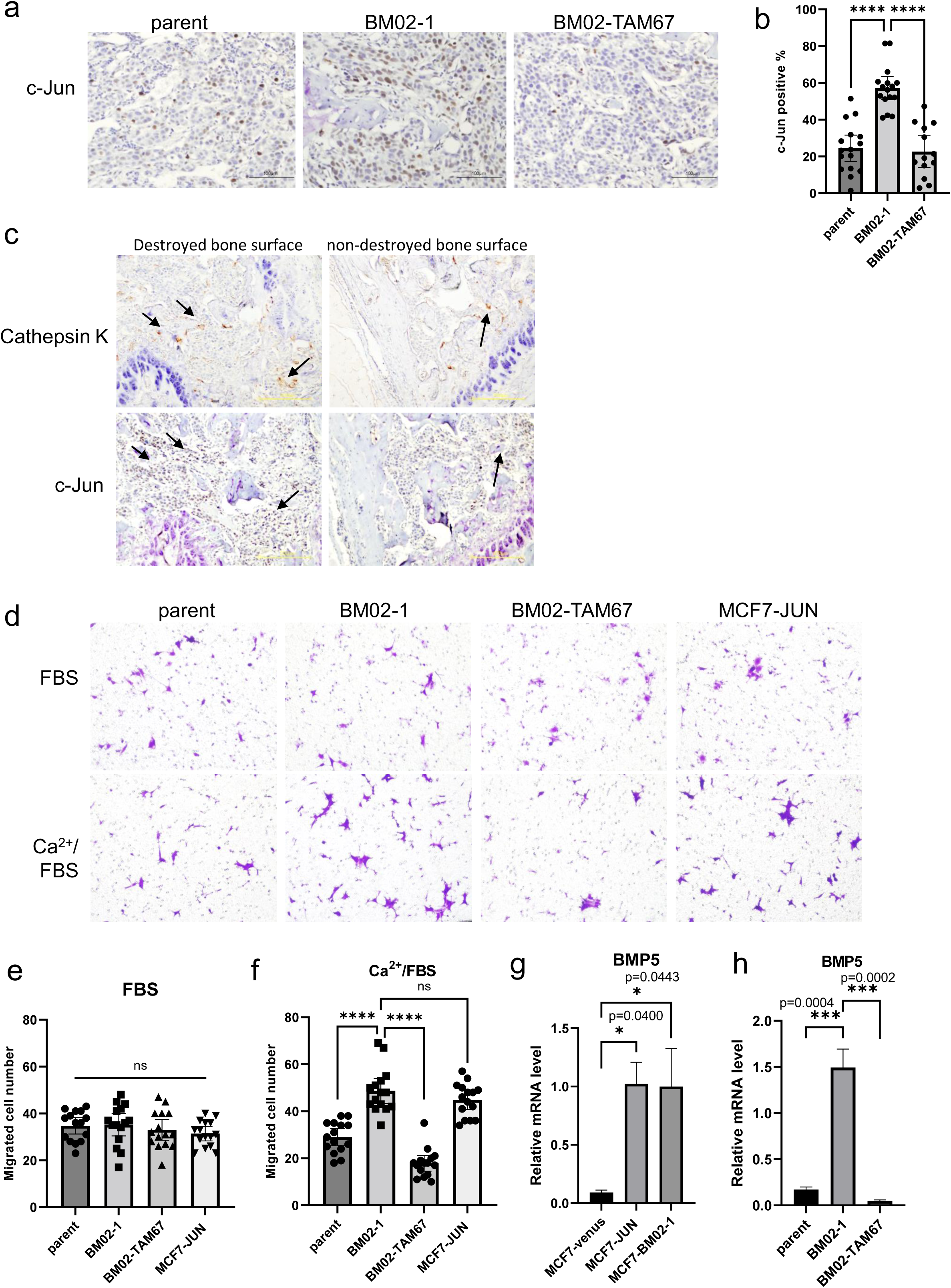
Regulation of a vicious cycle by c-Jun between the bone microenvironment and tumor cells. **a)** The IHC staining with anti-c-Juno metastatic lesions. **b)** Statistical summary of c-Jun histological expression. The data are presented as the mean ± 95%CI. The significance was calculated by one-way ANOVA with Dunnet’s test. **c)** The IHC staining of cathepsin K and c-Jun of the same bone section under ×10 magnification. The black arrow indicates the cathepsin K-positive osteoclasts or the tumor cells with highly expressed c-Jun. **d, e, f)** The result of trans-well migration induced by Ca^2+^. **d** Photos of stained migrated cells. One-way ANOVA with Dunnet’s test analysis of the data from two independent trans-well experiments. **e** Migrated cells under the control (FBS only) condition. **f** Migrated cells under the 2.5 mM Ca^2+^ condition. The data are presented as the mean ± 95%CI. **g, h)** RT-PCR data showing the relative mRNA level of BMP5 in the MCF7-parent, MCF7-JUN, MCF7-BM02-1, and BM02-TAM67. The statistical analysis was performed by one-way ANOVA with Dunnet’s test. The data are presented as the mean ± SEM.

Our data demonstrated that c-Jun induced a more invasive phenotype of luminal breast cancer cells. To analyze the downstream signals of c-Jun, we performed RNA-seq analysis of the MCF7-parent, MCF7-BM02-1, and MCF7-TAM67 cell lines. The results showed that BM02-TAM67 and MCF7-parent shared some common c-Jun-regulated genes (Supplementary Fig. S2a, b). The gene signatures common between the TAM67 and parental cell lines were extracted as c-Jun regulated gene signatures, among which metastasis-related genes, such as matrix metalloproteinase (MMP)-21^23^ and latent transforming growth factor-β-binding protein (LTBP)2^24^, were upregulated by c-Jun. We performed Gene Ontology (GO) term enrichment analysis using the c-Jun-regulated gene signatures (Supplementary Fig. S2c). Genes that were involved in cell secretion and migration were mostly enriched. The NABA secreted factors and SMAD protein phosphorylation signals were two top upregulated signals by c-Jun (Table 1). Moreover, some of the c-Jun-regulated genes were related to the poor clinical prognosis of luminal A breast cancer patients (Supplementary Fig. S3a, b).

**Table 1.**
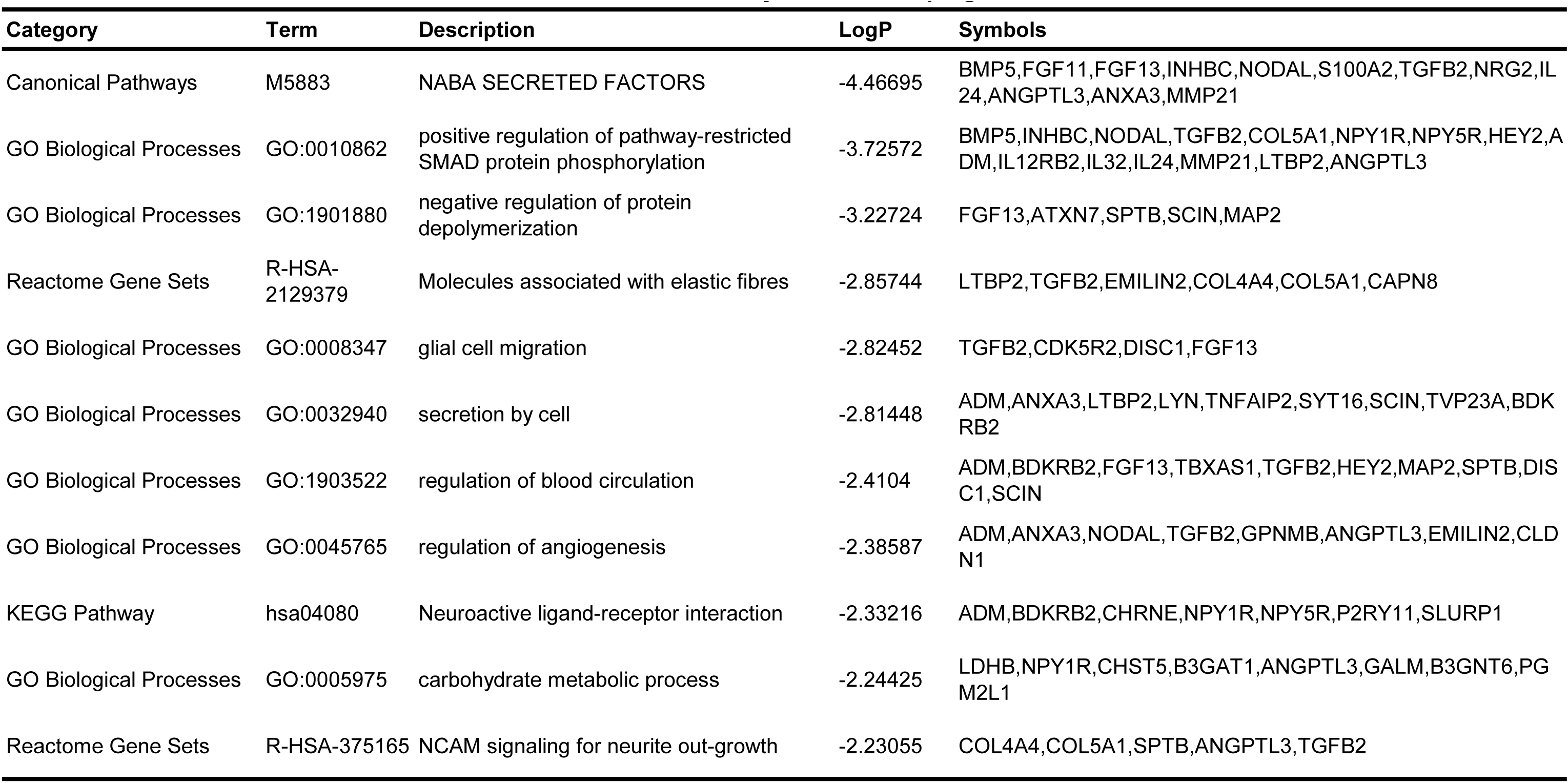
GO enrichment analysis of c-Jun upregulated DEGs

On the basis of the RNA-seq analysis results, we checked the downstream signals of c-Jun by real-time reverse transcription-polymerase chain reaction (RT-PCR). An osteoclast activator, bone morphogenetic protein 5 (BMP5), was revealed to be upregulated by c-Jun (Table 1, Fig. 5g, h). This finding was supported by previous research showing that BMP5 gene contained a DNA-binding site of AP-1 factor, 5′-TGATTCA-3′, in its promoter region^25^. It has been shown that BMP5 not only activates osteoclasts in a dose-dependent manner^26^, but also enhances osteoblast activity at higher concentrations. We thus performed a mineralization co-culture assay between osteoblasts and MCF7 cells. The results showed that MCF7-BM cells did not enhance osteoblast mineralization (Supplementary Fig. S4). These results suggested that the BMP5 released by MCF7-BM cells promoted osteoclast maturation but was insufficient to induce osteoblast activation.

### **c-**Jun inhibition as an effective therapy for luminal breast cancer bone metastasis

Our data supported the potential usefulness of c-Jun inhibitor therapy for luminal breast cancer bone metastasis. We applied a JNK inhibitor, JNK-IN-8, to inhibit the c-Jun activity in the MCF7-BM02-1 cell line. Complete inhibition of c-Jun phosphorylation was achieved at 2.5 μM JNK-IN-8 after 2 hours of incubation (Supplementary Fig. S5a, b). Treatment with JNK-IN-8 did not lead to cell death or significantly affect cell proliferation (Fig. 6a), but it did inhibit cell migration (Fig. 6b, c), transforming ability (Fig. 6d, e, f), and tumor stemness of BM02 cells (Fig. 6g, h). These results were consistent with those of BM02-TAM67 cells. We also found that JNK-IN-8 used in a co-culture assay with osteoclasts and MCF7-BM cells inhibited osteoclast activity with or without the participation of tumor cells (Supplementary Fig. S5c, d). On the basis of the *in vitro* result, JNK-IN-8 injections were applied to MCF-BM02 bone metastasis models (Supplementary Fig. S5e). The bone metastatic tumors treated with JNK-IN-8 were much smaller than those with DMSO vehicle control (Fig. 6i, j). Severe side effects were not observed during the treatment as shown by the weight curve of mice (Fig. 6k). The bone metastatic lesion sizes were suppressed from the 7^th^ day of observation, and progression of the lesions was inhibited throughout the 28-day observation period (Fig. 6l, m). Our results demonstrated that JNK-IN-8 was an efficient treatment for bone metastasis in luminal breast cancer.

**Fig. 6.**
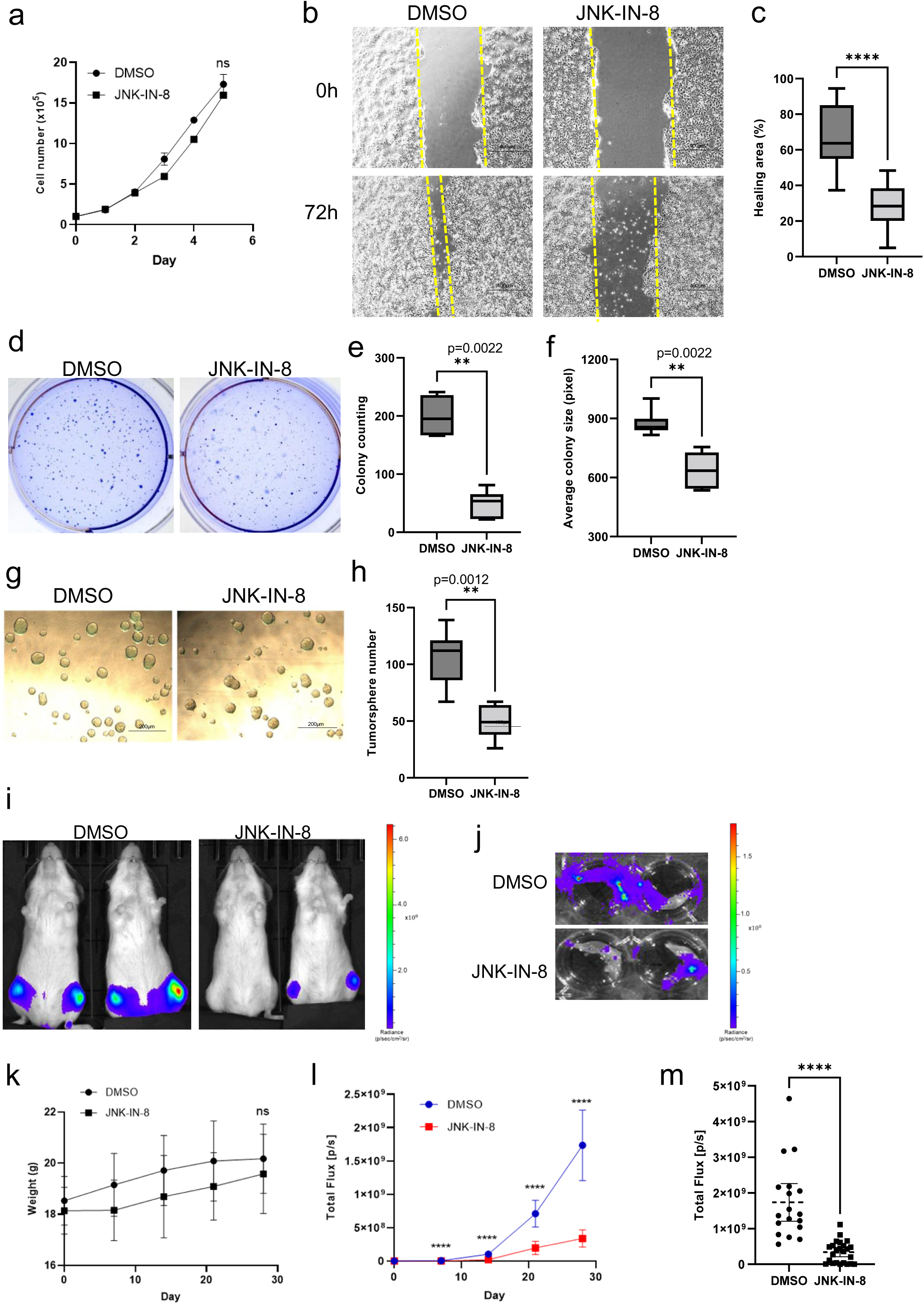
JNK-IN-8 inhibited bone metastatic lesions of luminal breast cancer. **a)** Growth curve of the BM02-1 cell line with JNK-IN-8 treatment. The data are presented as the mean ± SEM and analyzed by the *t*-test with Mann–Whitney analysis. **b, c)** Scratch assay of BM02-1 cells with JNK-IN-8 treatment. DMSO was added as a control. **b** Photos of the wound-healing process. **c** Statistical analysis of the healing area. The results were summarized from two independent experiments and analyzed by the *t*-test with Mann–Whitney analysis. **d, e, f)** Soft agar-formation assay of the BM02-1 cell line with JNK-IN-8 treatment. **d** Photos of crystal violet staining of colonies. **e** Statistical analysis of the number of colonies. **f** Statistical analysis of the average colony size by pixel. The results were summarized from two independent experiments and analyzed by the *t*-test with Mann–Whitney analysis. **g, h)** The tumorsphere assay of BM02-1 with JNK-IN-8 treatment. **g** Photos of tumorspheres at the 7^th^ day of culture. **h** Statistical analysis of the number of tumorspheres. The results were summarized from two independent experiments and analyzed by the *t*-test with Mann–Whitney analysis. **i, j)** Transplantation of BM02-1 to NOD/scid mouse through CAI. JNK-IN-8 was injected every 7 days. The transplantation was performed independently for three times (DMSO, n=9; JNK-IN-8, n=12). **i** *In vivo* imaging photos of a mouse with bone metastatic lesions. **j** *Ex vivo* imaging photos of legs that carried bone metastatic tumors. **k, l, m)** Statistical analysis of CAI. The data are presented as the mean ± 95%CI and analyzed by the *t*-test with Mann– Whitney analysis. **k** Weight change of the mouse during the transplantation. **l** Tumor growth through *in vivo* observation. **m** Summary of tumor size after 28 days of observation.

### Classification based on c-Jun signature genes of luminal breast cancer patients

To further illustrate the important role of c-Jun in breast cancer, we reanalyzed RNA-seq data of primary breast cancer patients from the The Cancer Genome Atlas (TCGA) and clustered them by using extracted c-Jun-regulated DEGs (Supplementary Fig. 2). The clinical patients were divided into three subgroups (Supplementary Fig. S6a, b). The luminal subtypes (Group A, Group B) were discriminated from basal breast cancer patients (Group C). Therefore, we extracted the luminal cohort from all breast cancer patients and clustered using c-Jun-regulated DEGs. The luminal subtype patients were classified into two subgroups, which showed significant differences in the clinical prognoses (Fig. 7a, b). These results suggest that c-Jun-regulated genes, namely, c-Jun signature genes, are useful as specific diagnostic markers of luminal breast cancer.

**Fig. 7.**
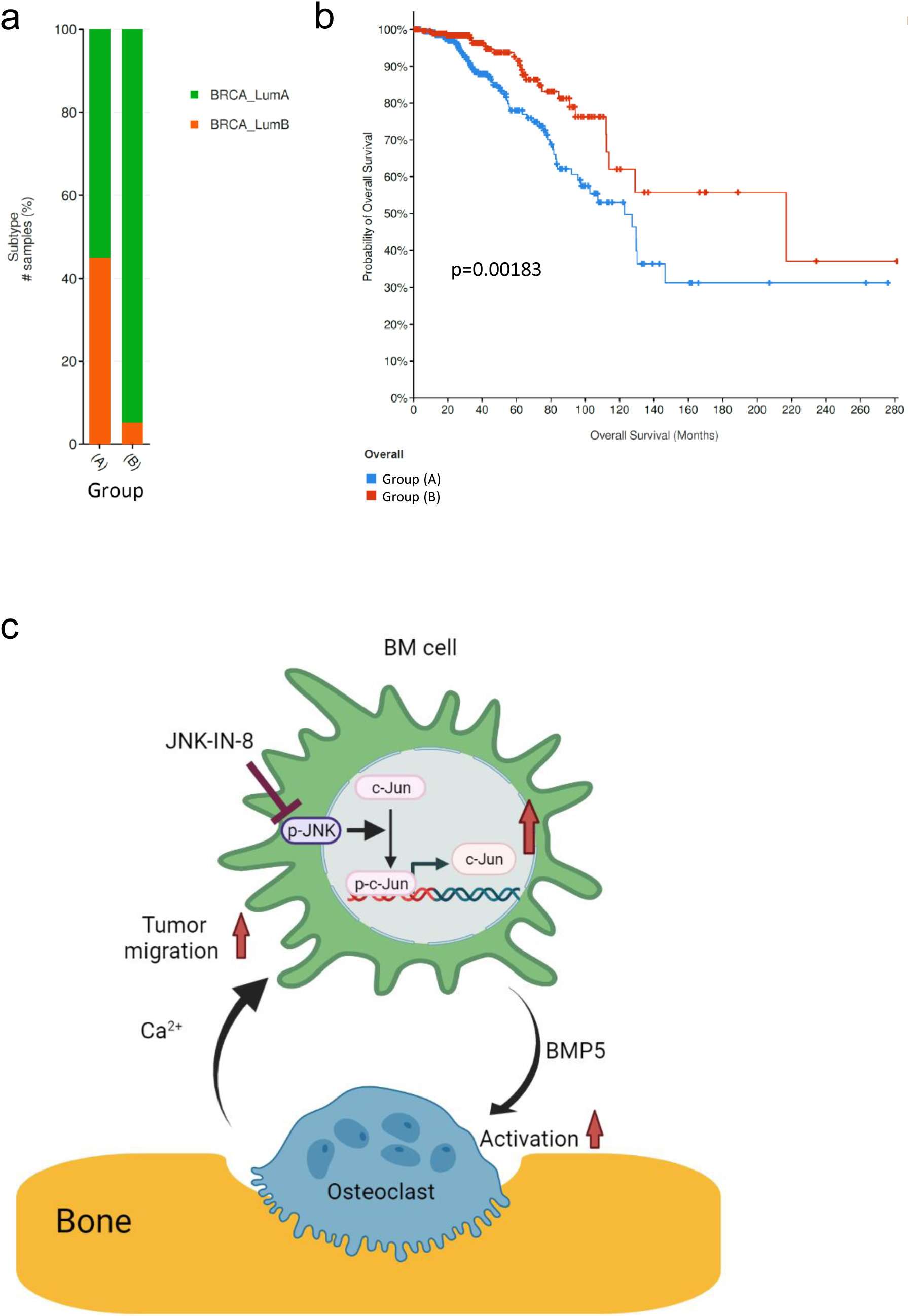
Clinical clustering of luminal breast cancer using c-Jun-regulated gene signatures. **a)** The luminal cohort was extracted from public data provided by TCGA and clustered using c-Jun-regulated DEGs (477 genes in total). The patients with luminal breast cancer were divided into two subgroups. **b)** Overall survival curves of the two subgroups. The *P* value was calculated by the *t*-test. **c)** The conclusion scheme of this study created with BioRender.com.

## Discussion

In this study, we used a highly bone metastatic cell line, MCF7-BM, to analyze the mechanism of bone metastasis in the luminal subtype of breast cancer. Our results indicated that c-Jun was a crucial factor in luminal breast cancer bone metastasis by promoting osteolytic metastasis of MCF7 in the bone microenvironment via Ca^2+^-induced cell migration and secretion of BMP5. Furthermore, inhibiting c-Jun by JNK-IN-8 suppressed the formation of bone metastasis lesions in luminal breast cancer (Fig. 7c).

Western blotting of signal transducers showed increases in the c-Jun protein at the post-translational level in MCF-BM cells. Previous studies have shown that various ubiquitin ligases, such as SCF^Fbw7^, HECT, and COP1^27–31^. However, our RNA-seq analysis showed that the RNA level of the reported c-Jun regulators did not change in BM02. Elucidation of the regulatory mechanism of c-Jun stability is still needed.

BMPs exhibit multiple functions during tumor progression. For example, BMP2 has been shown to promote bone metastasis of non-small cell lung cancer^32^, BMP4 has been shown to enhance bone metastasis of MDA-MB-231^33^, and BMP5 has been shown to activate osteoclasts in a dose-dependent manner^26^. Importantly, in clinical studies, BMP5 was found to be overexpressed in primary tumors of breast cancer and was associated with poor prognosis^34,35^. With no reports about the relationship between BMP5 and bone metastasis of breast cancer cells, our results suggested that high expression of c-Jun in MCF-BM cells activated osteoclast maturation by releasing BMP5 in the bone microenvironment.

In the present study, MCF7 cells migrated in fetal bovine serum (FBS) showed no difference in the trans-well migration assay, which seems to be inconsistent with the wound-healing assay result. It should be noticed that the wound-healing assay is used to evaluate collective-cell migration, whereas the trans-well assay is a single-cell migration assay^36,37^. In previous reports, MCF7 exhibited different migration phenotypes under stimulations or genetic mutations; for example, HCLS1-associated protein X-1 knockdown in MCF7 only affected wound-healing migration, not trans-well migration ability^38^, and staurosporine induced migration in single cells but inhibited collective MCF7 cell migration^39^. In the present study, compared with parental cells, MCF7-BM02 cells showed stronger collective-cell migration than single-cell migration ability, which was mediated by c-Jun protein expression. On the other hand, c-Jun enhanced single-cell migration of MCF7 to a high Ca^2+^ environment. A previous study showed that activation of the calcium-sensing receptor in a high Ca^2+^ environment had a chemoattractant effect on migration of highly bone metastatic breast cancer cells toward a Ca^2+^-rich environment^40^. The intracellular signals activated by calcium ions in estrogen receptor-positive breast cancer cell lines (including luminal A/B) have been shown to be mediated by the stromal interaction molecule 1/2 and calcium release-activated calcium-channel protein 3 axis^41^. However, the mRNA level of Orai3 showed no significant difference between MCF-BM cells and parental cells (data not shown). Clarification of the detailed mechanism of how BM cells enhance migration toward a Ca^2+^-rich environment is needed. In addition to Ca^2+^, the other factors released from destroyed bone structure need to be evaluated to prove the proposed vicious cycle.

A recent study identified the third-generation subtype classification of breast cancer patients based on treatment responses^42^. In the present study, we used c-Jun downstream signals to further classify the subtypes of luminal breast cancer and their clinical prognoses. Therefore, c-Jun downstream signals may help predict patients that can benefit from c-Jun inhibition therapy. We showed that several genes were related to the survival of luminal A patients (Supplementary Fig. S3). Among these genes, some have important roles in other breast cancer subtypes as follows: Bradykinin receptor B2 antagonist therapy has been shown to arrest growth and induce apoptosis in TNBC^43^, and downregulation of 5′-methylthioadenosine phosphorylase has been shown to promote tumor growth and lung metastasis of TNBC^44^. These downstream signals can be used to further explore and identify qualifying biomarkers of luminal breast cancer.

Since c-Jun exhibited multiple functions during bone metastasis, its inhibition is a potential therapy. JNK-IN-8 is an irreversible c-Jun N-terminal specific inhibitor^45^ that can increase the survival of MDA-MB-231 xenograft models with combined JNK-IN-8/lapatinib treatment, and its inhibition toward AP-1 factor has previously been shown to contribute to several cancer treatments, such as resistant basal cell carcinoma, paclitaxel (PTX)-resistant MCF-7 tumors, and lung metastasis of immunosuppressive TNBC tumors^46–49^. In the present study, although JNK-IN-8 did not suppress cell proliferation of MCF-BM *in vitro*, it significantly inhibited cell migration, colony-forming ability, and tumor stemness of MCF-BM cells. Additionally, c-Jun signaling has been shown to have a crucial role in RANKL-regulated osteoclast differentiation^50,51^. We consistently observed by TRAcP assay that JNK-IN-8 inhibited osteoclast differentiation (Supplementary Fig. S5c, d). These unique phenotypes of JNK-IN-8 enable effective suppression of bone metastasis through a dual effect on both tumor cells and the bone microenvironment. In models of JNK knockout mice, JNK knockout was observed to induce behavioral changes, resistance against toxicity-induced brain neuron cell death, and defection of T-cell differentiation^52–54^. Although JNK-IN-8 hardly passes through the blood–brain barrier, long-term observation is required to evaluate its effects on bone and immune and neural homeostasis.

In conclusion, our results suggested a novel model in which c-Jun leads to a potential vicious cycle in the bone microenvironment and promotes bone metastasis of the luminal subtype of breast cancer. These results also indicated the therapeutic possibility of JNK-IN-8. Further pre-clinical studies are necessary to evaluate the potential of the JNK–Jun axis as a therapeutic target of bone metastasis in luminal breast cancer.

## Material and methods

### Cell culture

Culturing of MCF7 cells (IDAC, Tohoku University, Miyagi, Japan) and their derivatives was performed in Roswell Park Memorial Institute (RPMI) (Fujifilm Wako, Osaka, Japan) supplemented with 10% FBS, 100 U/mL penicillin G, and 100 µg/mL streptomycin. Bone marrow cells were cultured in minimum essential Eagle medium, alpha modification (α-MEM; Life Technology, Carlsbad, CA, USA) supplemented with 10% FBS, 100 U/mL penicillin G, and 100 µg/mL streptomycin. All cell lines were incubated at 37°C in a 5% CO_2_ incubator.

### Vector construction

Human c-Jun sequence was amplificated using the cDNA of MCF7-BM02-1 and then transferred to pMXd3-PEF1-IRES-Puro vector using an In-Fusion Advantage PCR Cloning Kit (Clontech Laboratories Inc., CA, USA). The sequence encoding the transactivation domain of c-Jun in pMXs-Jun-IH was kindly provided by Dr. S. Watanabe (Fukushima Medical University) and was deleted by PCR using the two primers: 5′-CATGACTAGCCAGAACACGCTGCCCAGCGTCAC-3′ and 5′-TTCTGGCTAGTCATGGTTCTATCTCCTTCGAAG-3′. The resulting Jun-deletion mutant, TAM67, was transferred to the pMXd3-PEF1-IRES-Puro vector using an In-Fusion Advantage PCR Cloning Kit. The sequences of pMXd3-PEF1-TAM67-IRES-Puro and pMXd3-PEF1-JUN-IRES-Puro were confirmed by DNA sequencing.

### Establishment of MCF7-BM02-TAM67 and MCF7-JUN cell lines

MCF7-mSlc7a1-luc2 cells were previously established from MCF7 cells by infection with lentivirus vector (pLenti-Pubc-mSlc7a1-IRES-HygR) to express the ecotropic receptor^10,55^. The MCF7-BM02-1 cells were infected with retrovirus vector (pMXd3-PEF1-TAM67-IRES-Puro) and selected by puromycin to establish MCF7-BM02-TAM67 cell line, followed by the transfection protocol established in previous stud^55,56^. By applying the same infection method, MCF7-parent cells were infected with retrovirus vector (pMXd3-PEF1-JUN-IRES-Puro) and selected by puromycin to establish the MCF7-JUN cell line.

### Quantitative real-time reverse transcription-polymerase chain reaction (real-time RT-PCR)

Quantitative real-time RT-PCR was performed using THUNDERBIRD^®^ (TOYOBO, Osaka, Japan) as previously described^57^. The following primers were used: (c-Jun forward: 5′-GAGCTGGAGCGCCTGATAAT-3′, c-jun reverse: 5′-CCCTCCTGCTCATCTGTCAC-3′; BMP5 forward: 5′-CGTGAGAGCAGCCAACAAAC-3′, BMP5 reverse: 5′-TGGTGCTATAATCCAGTCCTGC-3′).

### Western blotting and antibodies

Western blotting was performed as described in a previous protocol^57^. Hypotonic buffer (10 mM HEPES-KOH (pH 7.9), 1.5 mM MgCl2, 10 mM KCl, protease inhibitor cocktail, and 0.15 mM DTT) was added in addition to fractionate nuclear lysates and cytoplasmic lysates. For the JNK inhibition experiment, 0.1, 0.5, 1, or 2.5 μM JNK-IN-8 (Cat no: S490, Selleck, Houston, TX) or 2.5 μL of DMSO (Fujifilm Wako, Osaka, Japan) was added to MCF7-BM02-1 cells and cultured at 37°C in a 5% CO_2_ incubator for 1 or 2 hours before collecting the cell lysate. The primary antibodies used in this study were as follows: c-Jun, #9165, Cell Signaling Technology, Inc. (CST), Danvers, MA; p-c-Jun (Ser73), #3270, CST; c-Jun (60A8), #9165S, CST; p-SAPK/JNK (T183/Y185), #9255S, CST; SAPK/JNK, #9252S, CST; c-Jun (D) (TAM67), sc-44, Santa Cruz Biotechnology, Inc., Santa Cruz, CA; Lamin A/C (4C11), #4777S, CST; c-Fos (E8), sc-166940, Santa Cruz; p-c-Fos, sc-81485, Santa Cruz; α-tubulin, 013-25033, Wako Pure Chemicals Industries, Osaka, Japan. All primary antibodies were diluted 1:1000 before using. The detailed dilution and second antibody conditions are listed in the extended data.

### Animal studies of TAM67 bone metastasis

Intra-CAIs of 1.0 × 10^6^ cells/100 µL phosphate-buffered saline (PBS) of MCF7-parent (control), MCF7-BM02-1 or MCF7-BM02-TAM67 were made to transplant the cells into 6-week-old female NOD.CB-17-Prkdc<SCID>/J mice (NOD/scid, Charles River Japan, Inc., Kanagawa, Japan)^56,58^. The mice were anesthetized with 2.5% isoflurane (Fujifilm Wako) during transplantation and bioluminescence detection. Bone metastasis was monitored for bioluminescence using an IVIS Lumina XRMS In Vivo Imaging System (PerkinElmer, Waltham, MA, USA) every 7 days. Each mouse was intraperitoneally injected with 3 mg D-luciferin (Gold Biotechnology Inc., St. Louis, MO, USA) in 200 µl PBS, and the bioluminescence was measured with binning and F/stop ranges suited to each bioluminescence level. The bones from legs were harvested from the mice after 28 days. The *ex vivo* observation of the bone lesion was performed using IVIS-XRMS with 1.5 mg D-luciferin in 1 ml PBS.

### Animal study of JNK inhibition of bone metastasis

A 20-mg/kg solution of JNK-IN-8 was prepared in the solvent suggested by the manufacturer (Selleck, https://www.selleck.co.jp/products/jnk-in-8.html). DMSO was used as a vehicle control in the same solvent. Then, 1.0 × 10^6^ cells/100 µL PBS of MCF7-BM02-1 was transplanted into 6-week-old female NOD.CB-17-Prkdc<SCID>/J mice by CAI. The drug was injected 4 hours after injection of the cells and repeated every week. The *in vivo* and *ex vivo* observations were performed as described in the previous paragraph.

### Histological analysis of TAM67 bone metastasis lesions

The forelimb and hindlimb samples with bone metastasis lesions were fixed with 4% paraformaldehyde (PFA, Fujifilm Wako)/PBS solution at 4°C overnight twice. After fixation, the forelimb and hindlimb samples were decalcified using 0.5 M EDTA (pH 7.0) for 1 month. Hematoxylin-eosin (H&E) staining was then performed as previously described^10^. Immunohistochemistry (IHC) staining was performed on each sample using anti-cathepsin K (1:100, Cat. No. ab19027; Abcam Japan, Tokyo, Japan), anti-Ki-67 (1:100, Cat. No. ab16667; Abcam Japan), and anti-c-Jun (1:600, 9165; CST, Danvers, MA). The IHC staining of cathepsin K was performed as previously described^10^. The IHC staining of c-Jun and ki-67 was performed using a VECTASTAIN Universal Elite ABC Kit (Vector Laboratories, USA). The sections were stained with a DAB Substrate Kit (Vector Laboratories) followed by dehydration and mounting. The number of ki-67- or c-Jun-positive cells was quantified using ImageJ plugin’s “Color deconvolution2.”

### Growth curve of MCF7 cell lines

To evaluate the growth curve, 2 × 10^4^ MCF7 cells were seeded into each cell of a 12-well plate. The cell number was counted every 24 hours for 6 days. For JNK-IN-8 inhibition, 1 × 10^5^ cells per well of MCF7-BM02-1 were seeded into 12-well plates with 2.5 μM JNK-IN-8 or DMSO treatment. The culture medium was refreshed every 2 days, and the cell number was counted every 24 hours for 5 days.

### TRAcP assay co-cultured with cancer cells

Bone marrow cells were obtained from the femurs of 6−8-week-old BALB/cCrSIc female mice (SLC, Shizuoka, Japan) and cultured in α-MEM supplemented with 50 ng/ml M-CSF. Human macrophage colony-stimulating factor (M-CSF)-dependent macrophages were collected after 72 hours, seeded into a 96-well plate at 1.5 × 10^4^ cells/well, and allowed to attach overnight. The medium was then replaced with 150 μl of culture medium containing 50 ng/ml human M-CSF and 100 ng/ml recombinant receptor activator of NF-κB ligand (GST-mRANKL) containing 300 cells/well MCF7 cells. For the JNK inhibition experiment, 2.5 μM JNK-IN-8 or DMSO was added, and the mixture was then cultured for 96 hours until mature osteoclasts formed. The M-CSF, GST-RANKL, and TRAcP staining protocol was generated and described previously^10^. Stained osteoclasts with more than two nuclei were counted as mature osteoclasts.

### Soft agar-formation assay

A 2.4% agarose gel was made from SeaPlaque^®^GTG^®^ Agarose (Lonza, Basel, Swiss) and mixed with 1× RPMI and 2×RPMI (Nissui, Tokyo, Japan) in the proportion of 1.2:1:1 as bottom gel. Then, the 2.4% agarose gel was mixed with 1×RPMI and 2×RPMI in the proportion of 1:3.328:1 as top gel. Next, 1.5 ml of bottom gel was added to the wells in the 6-well plate. A 1-ml aliquot of top gel containing 1 × 10^4^ cells of MCF7 was added to the top of the bottom gel. For the JNK inhibition experiment, 2.5 μM JNK-IN-8 or DMSO was added to the top gel. The plate was incubated at 37°C in a 5% CO_2_ incubator, and the colonies were fixed by adding 4% PFA/PBS after 3 weeks of culture. The colony was stained with 0.005% crystal violet/10% methanol. Image J software (free; https://imagej.nih.gov/ij/) was used to measure the colony number and size.

### Wound-healing assay

For this assay, 3 × 10^5^ cells/well of MCF7 were seeded into a 12-well plate and incubated overnight. The cell monolayer was scratched with a 200-μL bulk yellow tip (BM Equipment, Tokyo, Japan) to create the wound. The culture medium was replaced with 1% FBS RPMI and changed every day. For the JNK inhibition experiment, the culture medium was 10% FBS RPMI with 2.5 μM JNK-IN-8 or DMSO since the same cell line was used, and a low FBS concentration could not maintain cell survival with the inhibitor. Three individual photos were taken from at least three wells of each cell line. Image J was used to analyze the wound area. The healing area between the wound area was calculated at 0 and 72 hours of migration.

### Tumorsphere culture of MCF7 cells

Tumorsphere culture was performed using a MammoCult Human Medium kit (Veritas, Tokyo, Japan). First, 500 μL MammoCult basal medium with 10% MammoCult supplement, 8 mg/L Heparin (Sigma, St. Louis, MS), 0.96 mg/L hydrocortisone (Sigma, USA), and 0.5% methyl cellulose (Fuji Film Wako, Japan) was mixed with 1 × 10^4^ MCF7 cells and added to the wells of an ultralow-attachment 24-well plate (Corning, New York, USA). For the JNK inhibitor experiment, 2.5 μM JNK-IN-8 or DMSO was added. The plate was incubated at 37°C in a 5% CO_2_ incubator for 7 days. Tumorspheres with diameters > 100 μm were counted under a microscope.

### Trans-well cell migration assay

A trans-well with an 8.0-µm pore polycarbonate membrane in a 24-well plate (Corning, US) was used to perform the trans-well cell migration assay. Then, CaCl_2_ was added to the well (final concentration of Ca^2+^ is 2.5mM) with 10% FBS RPMI. Next, 5 × 10^4^ MCF7 cells were seeded on the insert in serum-free RPMI medium. After 24 hours of incubation, the migrated cells were fixed with 4% paraformaldehyde (PFA)/PBS and stained with 0.1% crystal violet/10% MeOH. Five photos were taken through a microscope at x10 magnification, and the migrated cell number was measured by Image J^59^.

### Mineralization assay co-cultured with tumor cells

The pre-osteoblasts were collected from mouse calvaria of newborn BALB/cCrSlc pups (0–3 days after birth), and cultured in α-MEM for 2–3 days until full confluence. Mature osteoblasts were transferred into a 6-well plate at 8 × 10^5^ cell/well and cultured in α-MEM, 50 mg/ml L-ascorbate (TCI, Tokyo, Japan), and 2 mM β-glycerophosphate (Cayman Chemical Company, Michigan, US) (os-α-MEM) to provide an osteogenic supply. After 7 days of incubation for mineralization, MCF7 cells were added at 3000 cells/well and co-cultured for an additional 10 days with the medium replaced every 2–3 days^60^. Culture medium was removed from each well. The monolayer was fixed in 4% PFA/PBS for 10 minutes at room temperature. Alizarin red S solution (Sigma-Aldrich, USA) was added and incubated at room temperature for 20 minutes with gentle shaking. The cells were then washed five times with MilliQ water. Then, 10% acetic acid was added and incubated at room temperature for 30 minutes with shaking to dissolve the pigment. The supernatant was collected into a new 1.5-mL tube, and the acidic solution was neutralized with 10% ammonium hydroxide. The absorbance at 450 nm was measured by a TriStar² LB 942 microplate reader (Berthold Technologies, Bad Wildbad, Deutschland).

### RNA-sequencing analysis

The RNA-sequencing was performed by GENEWIZ’s RNA-sequencing service (Azenta Life Sciences, USA). The genome was mapped and assembled using *Homo sapiens* reference genes GRCh38.p13 provided from Ensembl^61^ by using HISAT2^62^ and String Tie^63^ methods. Genes from the MCF7-parent and BM02-TAM67 were compared with those from BM02-1. The DEGs were calculated and standardized using edgeR^64^. The gene signatures were extracted followed by statistical analysis: log fold change (FC) > 1.0 or < −1.0, *p* < 0.05, FDR < 0.25. The heatmap was drawn using the “pheatmap” package of R version 4.1.2, and the clustering method was “wardD2.” The GO term enrichment analysis was performed using Metascape. A Venn diagram was drawn using the “ggplot2” package in R version 4.1.2.

The RNA-seq data of primary breast cancer patients were obtained from TCGA. The heatmap was drawn using the “pheatmap” package of R version 4.1.2, and the clustering method was “wardD2.” The overall survival curves of subgroups were generated by using cBioPortal^65^.

### Survival analysis

Survival analysis of c-Jun-regulated DEGs was performed using the Kaplan–Meier method for patients with luminal A breast cancer in the Molecular Taxonomy of Breast Cancer International Consortium (METABRIC) data set, as described previously^56^.

### Statistical analysis

One-way ANOVA with Dunnett’s test, Welch’s, and the Mann–Whitney *t*-test were performed using GraphPad Prism 9.00 (GraphPad Software, CA, USA). **** *p* < 0.0001, *** *p* < 0.001, ** *p* < 0.01, * *p* < 0.05, and ns (not significant) was assumed for *p* > 0.05. The log-rank test was performed using the “survminer” package in R version 4.1.2. Statistical significance for prognosis was defined as *p* < 0.05.

### Data availability

The study data are available from the corresponding author upon reasonable request. RNA-seq data were deposited into the NCBI GEO database (accession number: GSE205924).

### Code availability

The code for bioinformatic analysis has been available in the GEO database since Jun 14, 2022.

## Acknowledgments

We thank all laboratory members for the meaningful comments and discussion.

## Author contributions

YH performed the cell and animal experiments. YH and KN performed the bioinformatics analyses. YH and MF performed histological experiments. YH, YW, and MS performed the genetic experiments. KS and JN interpreted the data. YH wrote the manuscript. YH, KS and JN conceived and designed the study. All the authors reviewed and edited the manuscript.

## Ethical approval

The animal experiments were conducted under the approval of the ethics committee of Waseda University (2020-A067, 2021-A074).

---

**Supplementary Fig. S1.**
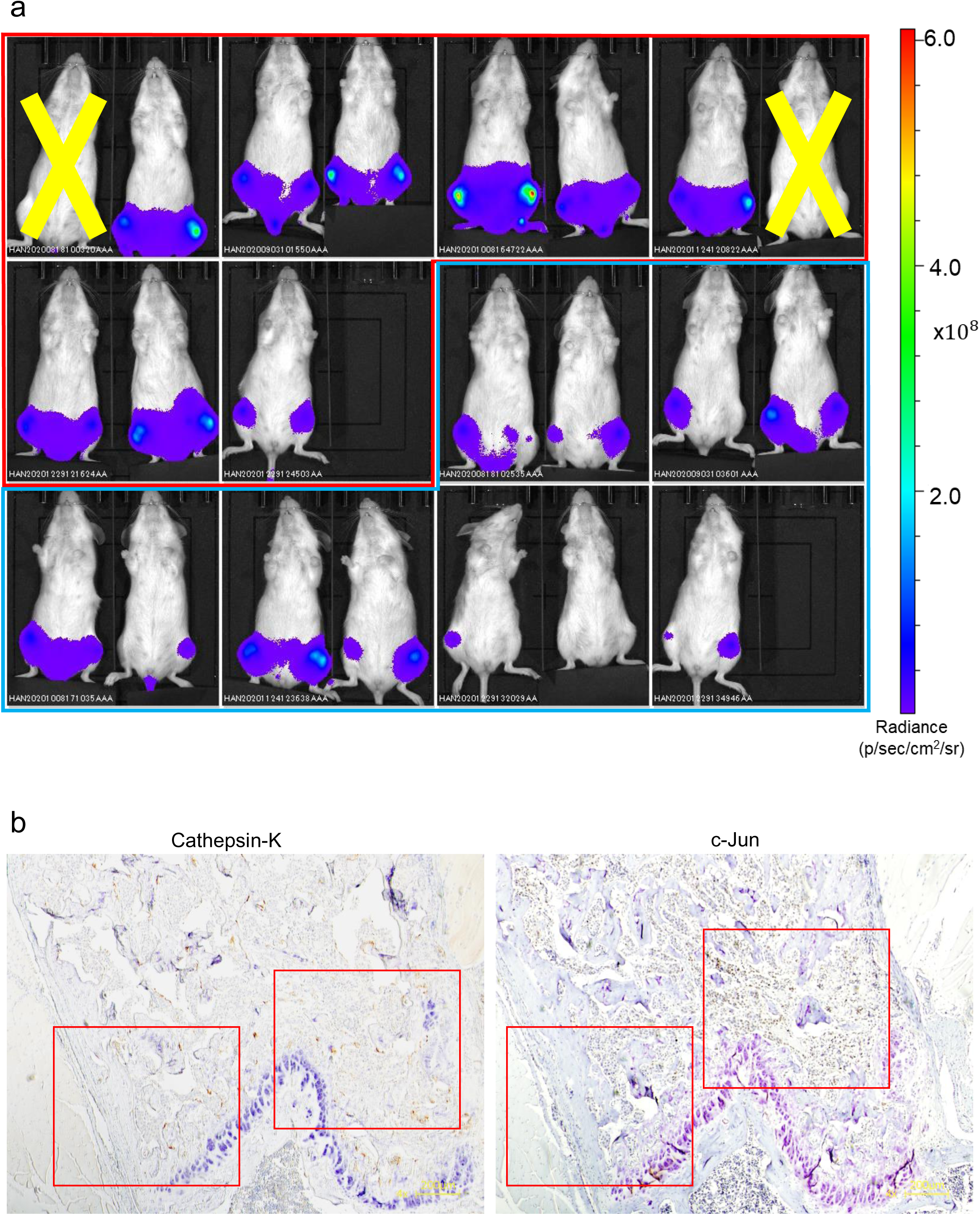
Evaluation of the bone metastasis potential of the TAM67 cell lines. **a)** The bone metastatic tumors of all transplanted mice at day 28 were observed and summarized. Bioluminescence was detected by IVIS. Red: MCF-BM02-1, n = 9. Blue: BM02-TAM67, n = 11. The mouse with a yellow cross sign failed transplantation. **b)** Zoomed IHC photos of cathepsin K and c-Jun of the same bone section at ×4 magnification. The red square indicates the ×10 magnification areas of Fig. 5c.

**Supplementary Fig. S2.**
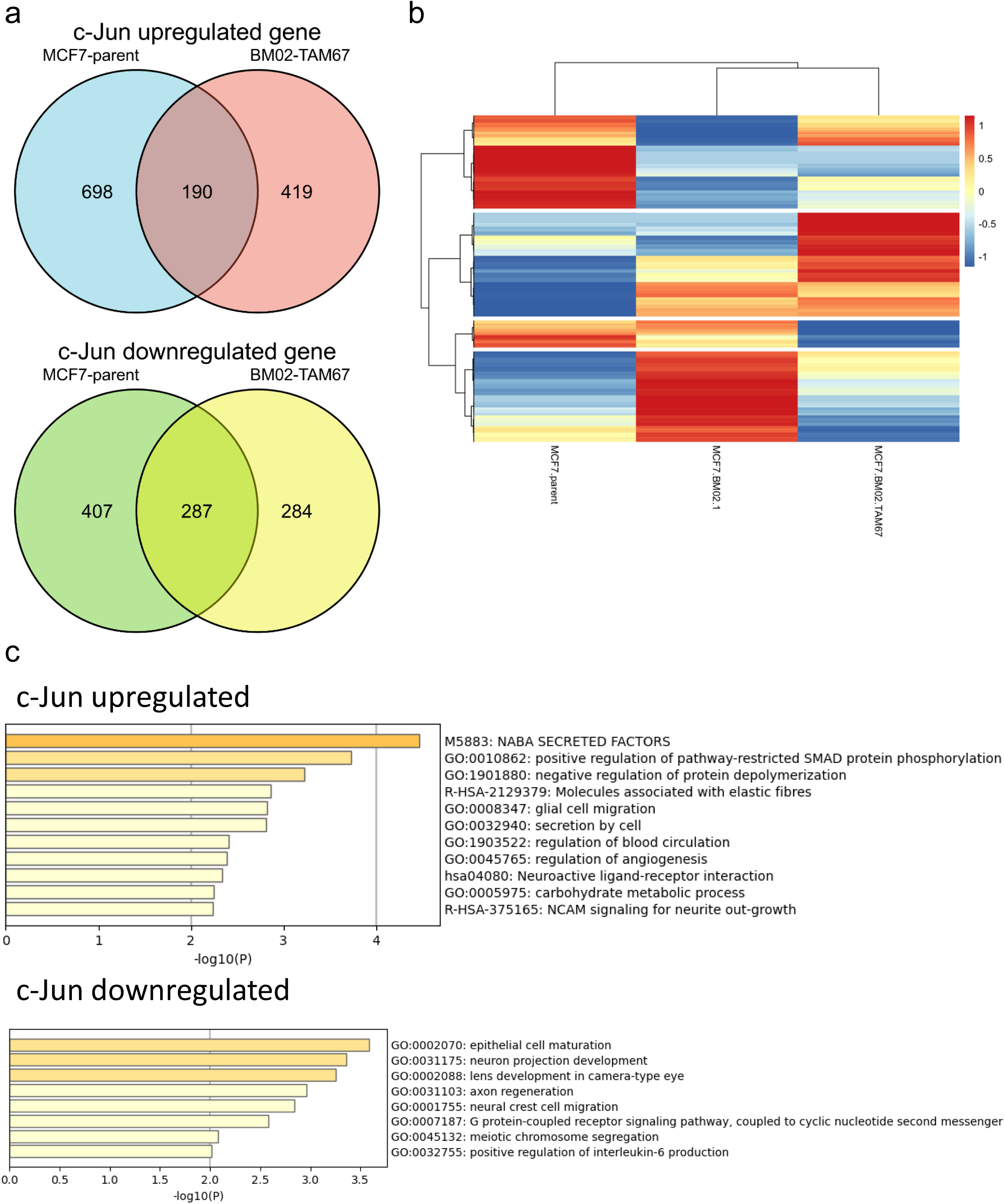
Extraction of c-Jun-regulated bone metastasis gene signatures from the MCF7 cell lines. **a)** The number of c-Jun-upregulated and -downregulated differential expression genes (DEGs) in the MCF7-parent and BM02-TAM67 cells compared with MCF7-BM02-1 cells were extracted under followed conditions: logFC ≥ 1.0 or ≤ −1.0, *p* < 0.05 and FDR < 0.25, and summarized in the Venn diagrams. The commonly shared gene signatures between the MCF7-parent and BM02-TAM67 were considered to be c-Jun-regulated DEGs. **b)** The extracted genes from the MCF7-parent and BM02-TAM67 cells were further analyzed by hierarchical clustering and are shown in a heatmap. **c)** The Gene Ontology (GO) enrichment analysis of c-Jun-upregulated and -downregulated DEGs.

**Supplementary Fig. S3.**
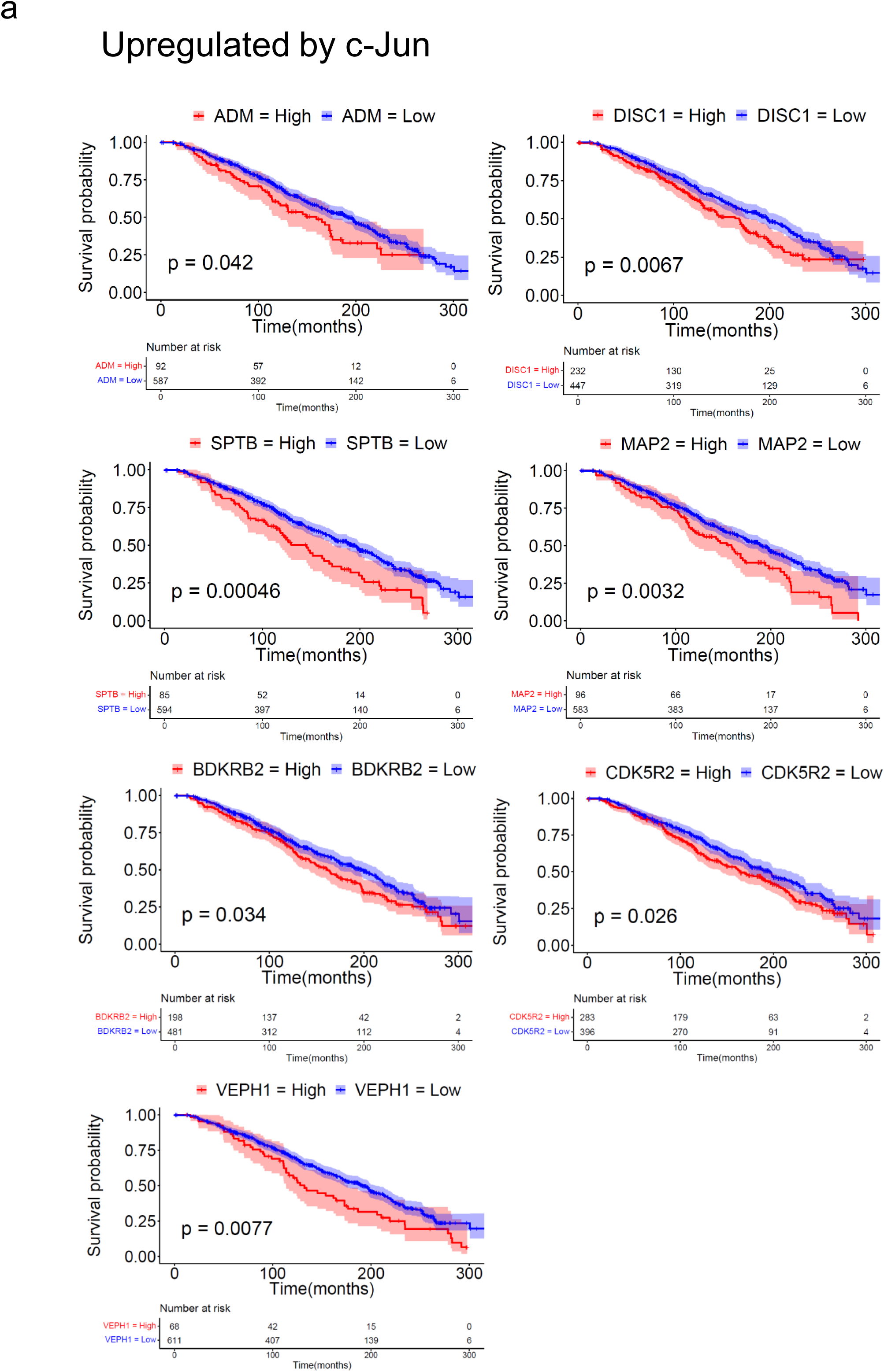
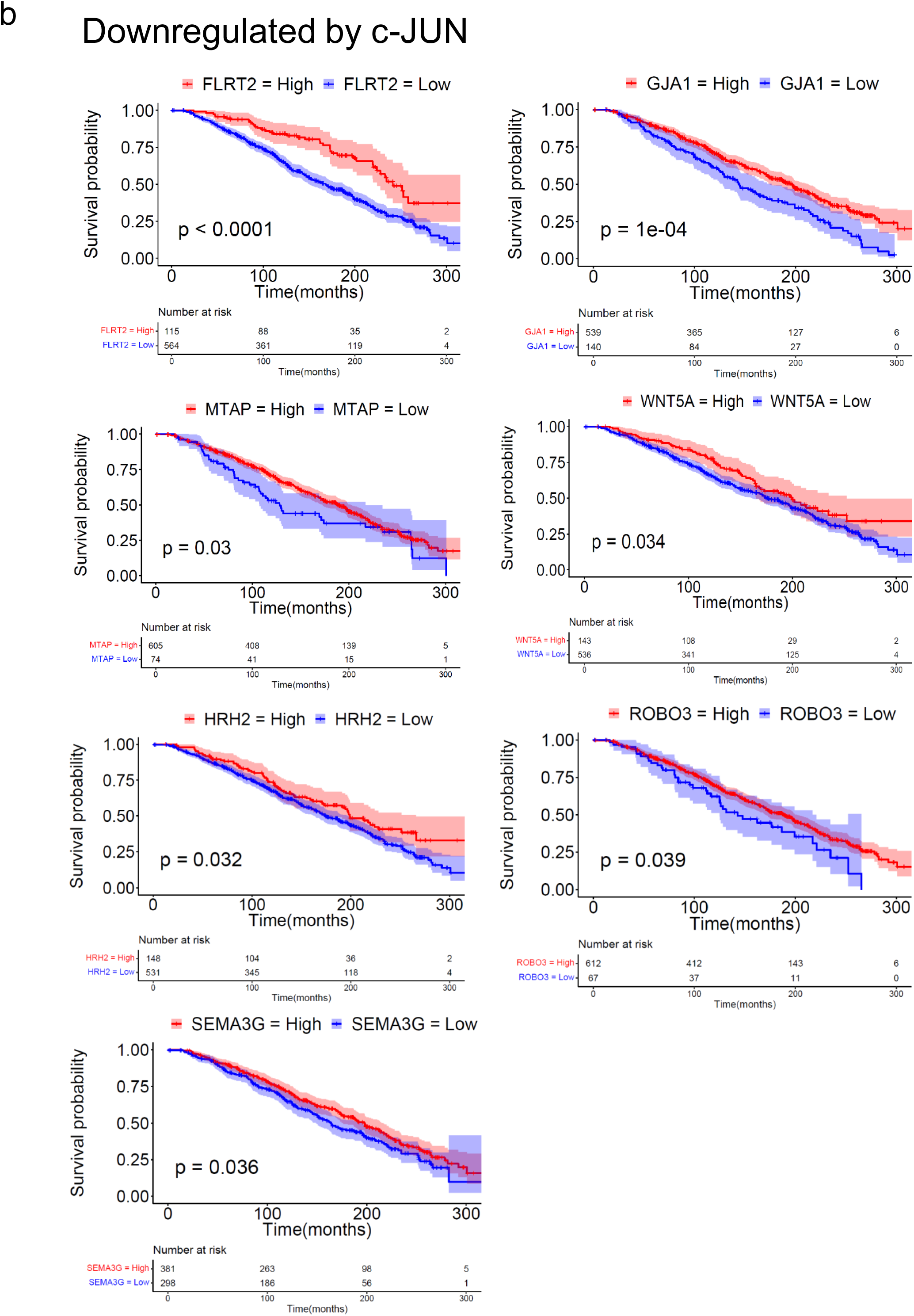
Clinical prognosis of c-Jun-regulated gene signatures. Survival analysis of **a)** c-Jun-upregulated and **b)** c-Jun-downregulated DEGs using the METABRIC data set. The colored region along the curve shows the 95% CI. The table at the bottom lists the number of patients with high or low gene expression in the luminal A subtype.

**Supplementary Fig. S4.**
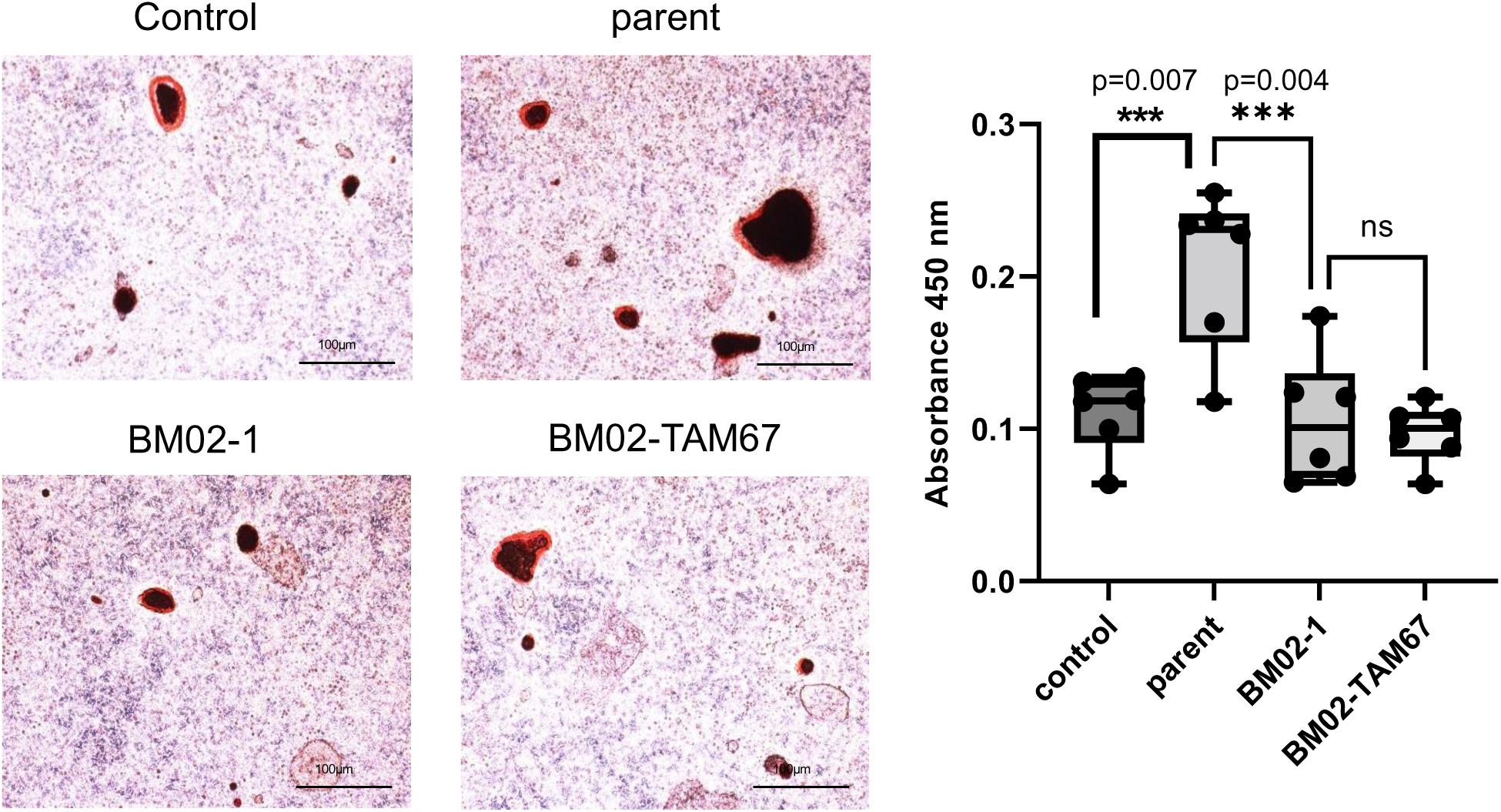
Mineralization assay co-cultured with MCF7 cells. The mineralization co-culture assay for MCF7 cells with osteoblasts. The control is the osteoblast culture without tumor cells. The calcium nodules formed by osteoblasts are stained a reddish color with Alizarin Red. The left figure shows the photos of the stained calcium nodules. The right figure shows the statistical analysis of the stained calcium nodules. The data were summarized from two independent experiments. The analysis was performed by one-way ANOVA with Dunnet’s test.

**Supplementary Fig. S5.**
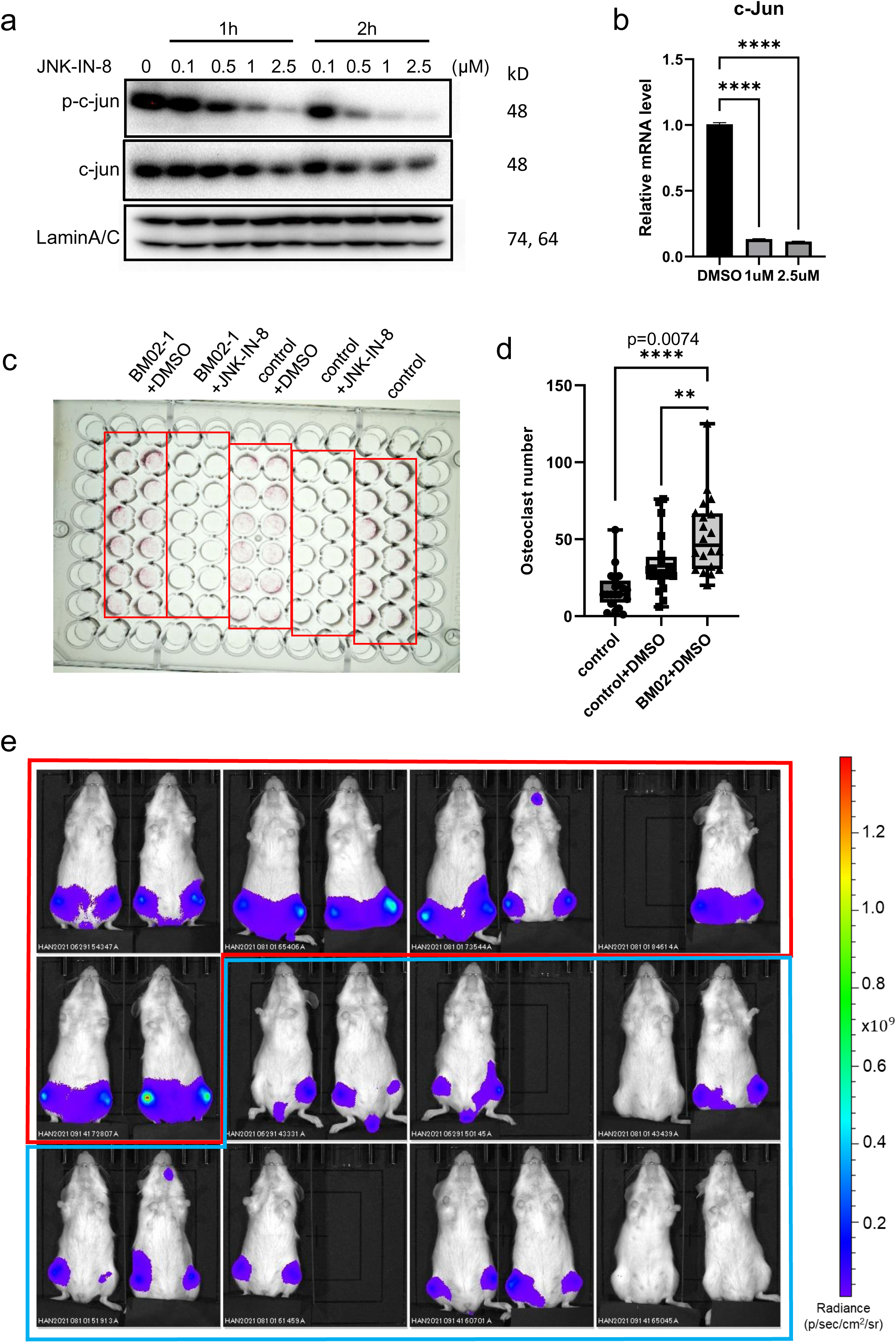
Evaluation of JNK-IN-8 inhibition of MCF7-BM cells. **a, b)** Evaluation of JNK-IN-8 inhibition of MCF7-BM02-1. **a** western blotting of c-Jun and phosphorylated c-Jun expression under a JNK-IN-8 gradient. **b** The mRNA level of c-Jun by RT-PCR. The data are presented as the mean ± SEM and analyzed by one-way ANOVA with Dunnet’s test. **c, d)** The TRAcP co-culture assay for BM02-1 cells and osteoclasts with or without JNK-IN-8 treatment. **c** Photo of a TRAcP-stained plate in which wells with additional JNK-IN-8 have completely lost mature osteoclasts. **d** Statistical analysis using one-way ANOVA with Dunnet’s test. The data were summarized from two independent experiments. **e)** Shown are the summarized photos of all bone metastatic tumors at day 28. The bioluminescence detected by IVIS. Red: DMSO (control), n = 9. Blue: JNK-IN-8, n = 12.

**Supplementary Fig. S6.**
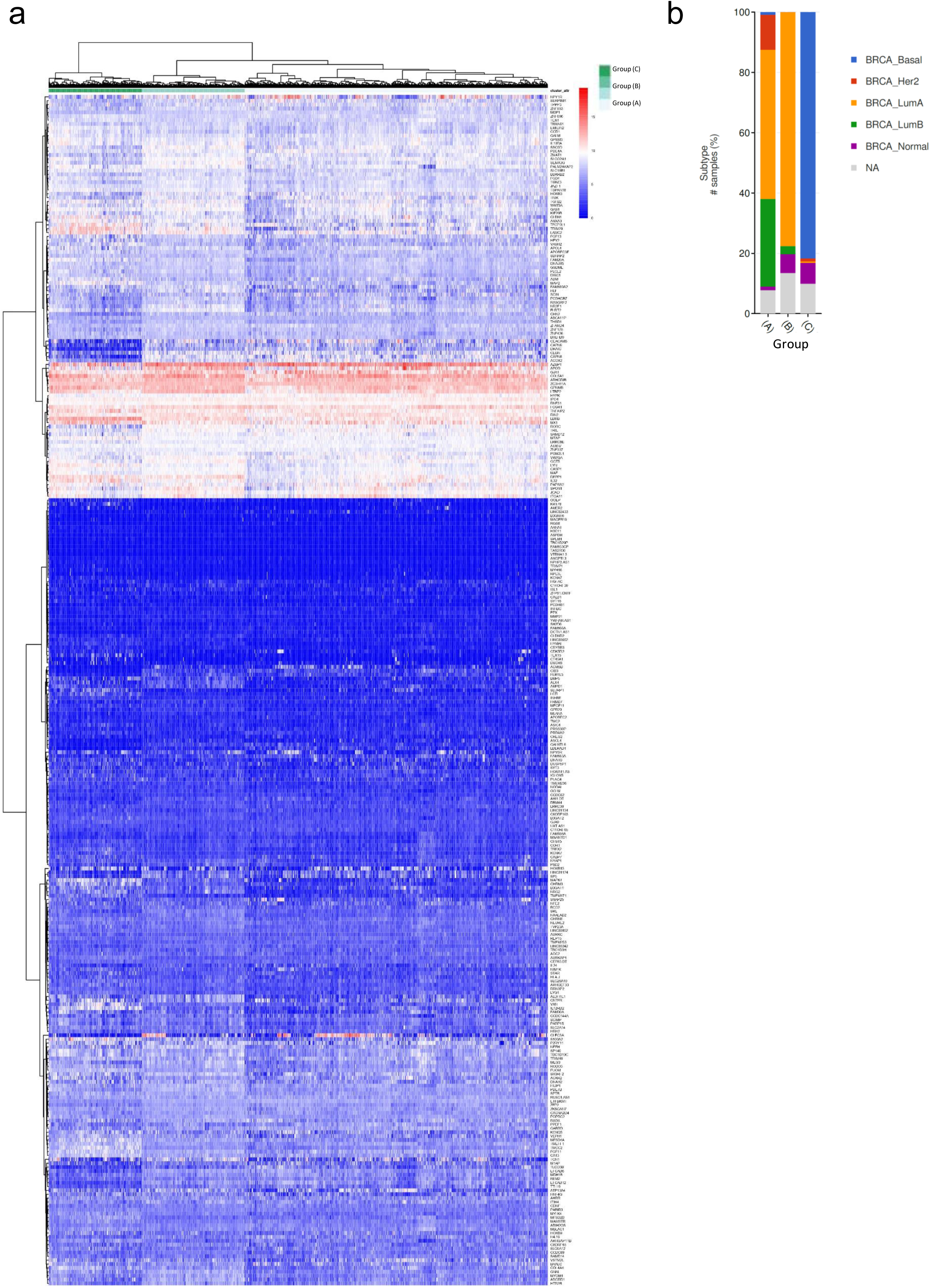
Clinical clustering of breast cancer subtypes using c-Jun-regulated gene signatures. **a)** The clinical data provided from TCGA clustered using c-Jun-regulated DEGs (477 genes in total). The patients with primary breast cancer were divided into three subgroups. **b)** Shown are the contents of each subgroup.

